# General finite-element framework of the Virtual Fields Method in Nonlinear Elasticity

**DOI:** 10.1101/2021.05.10.443225

**Authors:** Yue Mei, Jiahao Liu, Xu Guo, Brandon Zimmerman, Thao D. Nguyen, Stéphane Avril

## Abstract

This paper presents a method to derive the virtual fields for identifying constitutive model parameters using the Virtual Fields Method (VFM). The VFM is an approach to identify unknown constitutive parameters using deformation fields measured across a given volume of interest. The general principle for solving identification problems with the VFM is first to derive parametric stress field, where the stress components at any point depend on the unknown constitutive parameters, across the volume of interest from the measured deformation fields. Applying the principle of virtual work to the parametric stress fields, one can write scalar equations of the unknown parameters and solve the obtained system of equations to deduce the values of unknown parameters. However, no rules have been proposed to select the virtual fields in identification problems related to nonlinear elasticity and there are multiple strategies possible that can yield different results. In this work, we propose a systematic, robust and automatic approach to reconstruct the systems of scalar equations with the VFM. This approach is well suited to finite-element implementation and can be applied to any problem provided that full-field deformation data are available across a volume of interest. We also successfully demonstrate the feasibility of the novel approach by multiple numerical examples. Potential applications of the proposed approach are numerous in biomedical engineering where imaging techniques are commonly used to observe soft tissues and where alterations of material properties are markers of diseased states.

**List of symbols:** 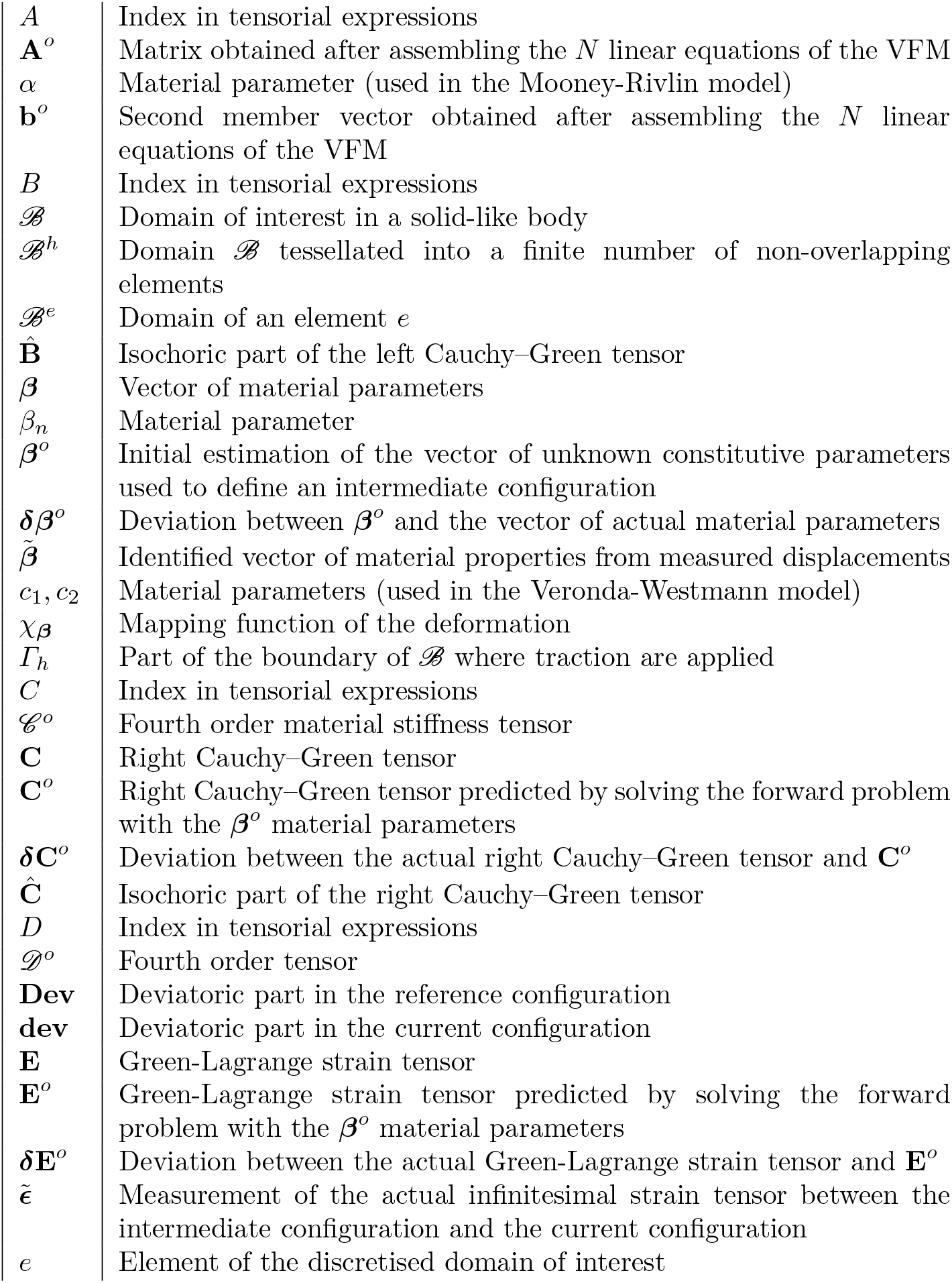

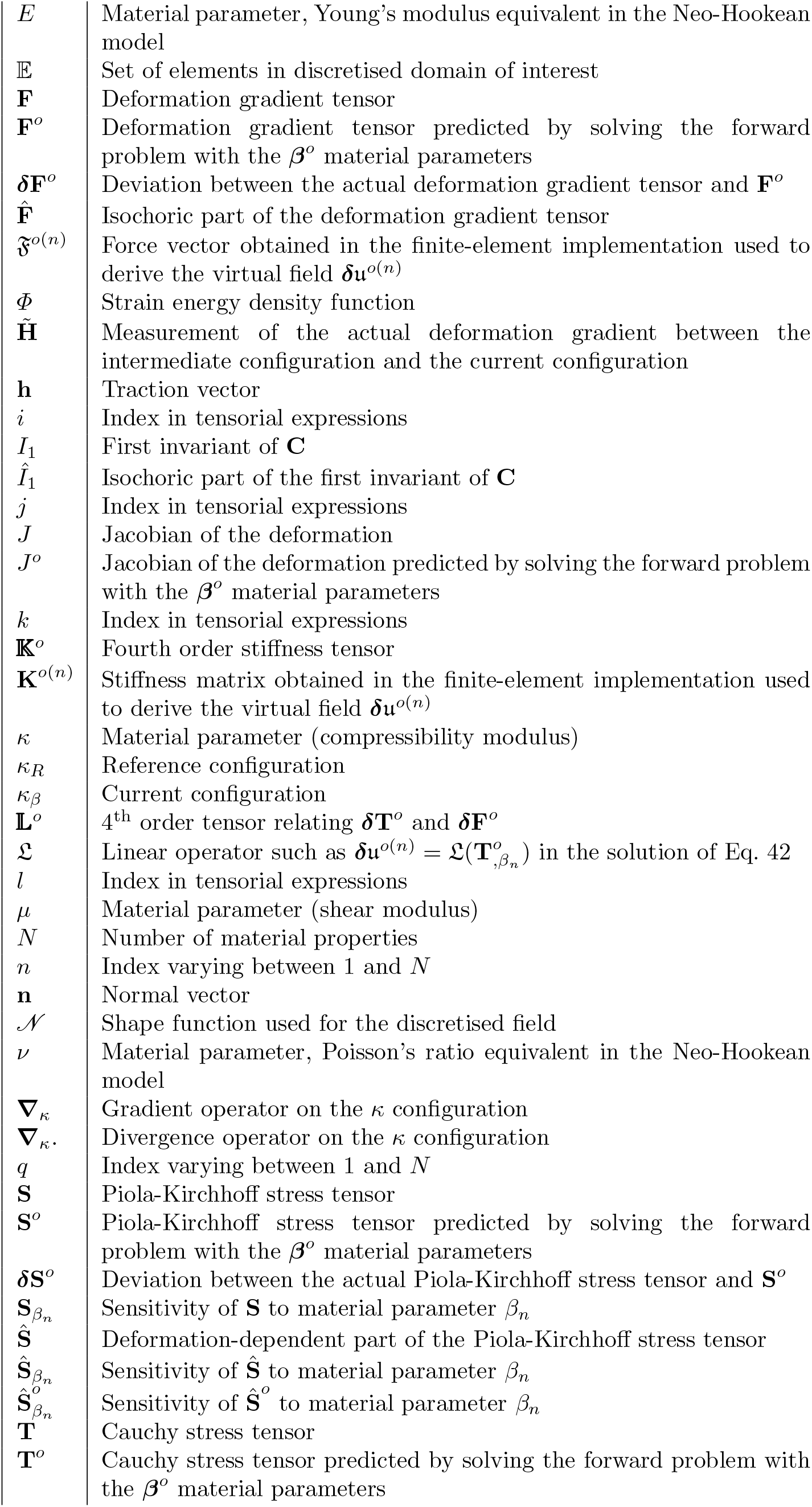

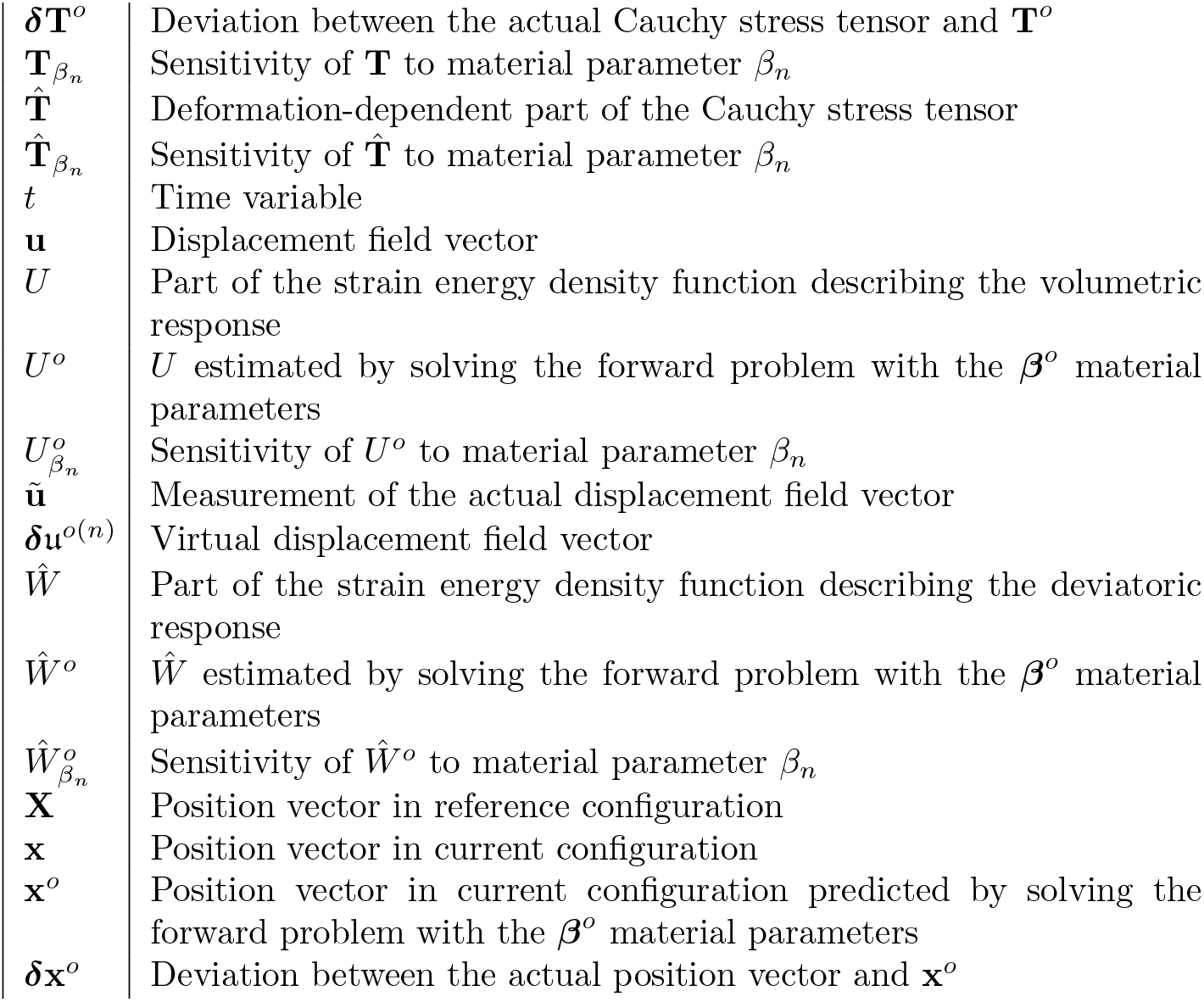

## 1 Introduction

One of the objectives in computational biomechanics is to make predictions: given for instance a complete and patient-specific reconstruction of the aorta and surrounding tissues, we can predict the deformations induced by a deployed stent graft during surgical repair of an aneurysm [1, 2, 3]. More generally, we could predict the deformations of the aorta under the action of any radial forces applied in an experiment. This problem of predicting the result of measurements is called the simulation problem, or the forward problem. The inverse problem consists of using the experimental measurements, such as the displacement field, to infer the values of the parameters that characterize the system, such as unknown material parameters [4, 5, 6, 7, 8, 9, 10, 11], unknown boundary conditions, or even sometimes the unknown initial geometry of the solid before the application of mechanical loading (load-free configuration in finite deformations [12, 13]).

A subcategory of inverse problems is made by identification problems [6, 14], where a finite number of unknown material parameters have to be recovered from experimental measurements. Such inverse problems can be solved by defining a cost function that estimates the absolute error between the model predictions and the measurements. The cost function is minimized in the least-squares sense. In general situations, the model is solved numerically using a finite-element model updating technique (FEMU) [6, 14, 15, 16, 17, 18, 19].

In the more and more common situations when full-field measurements are available [20, 21, 22, 7, 23, 24, 25], an alternative to FEMU is possible: the Virtual Fields Method (VFM), which has been shown to be more direct and robust with respect to unknown boundary conditions in these situations [26].

The general principle for solving identification problems with the VFM is first to derive parametric stress field across the volume of interest from the measured deformation field, stress components at any position for a given deformation depending on a number of unknown constitutive parameters. Applying the principle of virtual work to the parametric stress field gives scalar equations for the unknown parameters. The system of equations can be solved for the values of unknown parameters.

There are currently multiple strategies possible for the selection of virtual fields when the VFM is applied in nonlinear elasticity. A simple and efficient approach, well suited for tensile tests, is to define virtual displacements varying linearly from one boundary to the other. This approach was recently applied successfully in biaxial tension by Kazerooni et al [27] to identify the parameters of a Holzapfel model [28] for the skin. For other types of loading, for instance, Zhang et al. [23] introduced analytical expressions of virtual fields which were written in the cylindrical coordinate system using the *arctan* function. For inflation experiments, Bersi et al [29] used virtual fields that permitted to derive local equilibrium equations relating the intraluminal pressure and the wall tensile stresses. These virtual fields were valid for the very specific problem of inflation of thin-wall cylinder-like structures. The extension to thick-wall was solved in further papers [30, 25] by weighting locally the virtual fields with a Gaussian function, which was successfully centered at the middle of square patches in order to identify the distribution of material properties across the whole cylinder.

Although the approaches discussed above were all successful in solving specific identification problems, a general and unified VFM approach is still needed for the following reasons:

1. the expression of virtual fields used in most of previous studies [27, 30, 31] was well adapted to the 2D geometries such as membranes or shells but there is a range of samples or materials in which boundary conditions or geometries or both cannot be straightforwardly designed or chosen to be 2D. With the acquisition of fully volumetric deformation fields which is expanding for soft tissues [23, 24, 25, 32, 33, 34, 35], there is a need for a method that can provide virtual fields for any 3D geometry and virtual field.
2. In previous studies cited above, the VFM was used to identify the parameters of incompressible hyperelastic materials. The compressibility modulus was usually disregarded. To identify simultaneously the compressibility and shear moduli of a hyperelastic material, two independent virtual fields are needed in order to separate the hydrostatic and deviatoric contributions of the stress. This problem was solved a decade ago for linear elasticity [36, 37] but it remained an open question for nonlinear elasticity.

In this paper, we introduce an original approach addressing these essential questions. An application of this method to identifying material properties of the lamina cribrosa (LC) in the optic nerve head (ONH), using optical coherence tomography (OCT) imaging data, demonstrates the viability of the technique. The paper is organized as follows: We first elaborate the theoretical aspects of the proposed VFM approach in Section 2. Subsequently, several numerical examples and an application are presented to test the feasibility of the method in Section 3. We then discuss the results and the proposed approach in Section 4 and end with a conclusion in Section 5.

## 2 Materials and methods

### 2.1 Definition of the problem

A solid-like body, or part of it, denoted *ℬ*, is considered in a reference configuration *κ*_*R*_(*ℬ*). Under the application of loads, it is assumed that the solid has a nonlinear elastic response which is governed by a finite number *N* of material properties (defining local constitutive equations and their possible spatial variations).

These are denoted as a vector *β* = (*β*_1_, *β*_2_, …, *β*_*q*_, …, *β*_*n*_, …, *β*_*N*_).

At time *t*, after the application of traction **h**(*t*) on a part of the boundary of *B*, denoted by *Γ*_*h*_, *ℬ* undergoes a motion described by a mapping *χ*_***β***_, depending on material properties *β*, from a reference configuration *κ*_*R*_(*ℬ*) to a current configuration *κ*_***β***_(*ℬ*), such as

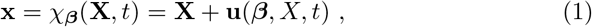

where **X** and **x** are position vectors relative to reference and current configurations and **u** is the displacement vector field. In the following, we introduce the deformation gradient

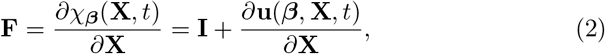

the right Cauchy–Green tensor

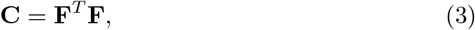

and the Jacobian

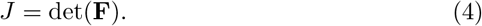

We also introduce the isochoric right Cauchy–Green tensor

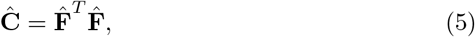

where we denote the isochoric part of the deformation gradient as

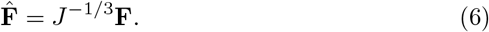

We assume that an experimental measurement of the displacement field **u**(**X**, *t*) is available across the solid at time *t*. The measured displacement field **ũ**(*t*) may be different than the actual displacement **u**(**X**, *t*) because of measurement noise.

The inverse / identification problem consists of finding the values of the approximate parameters 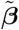 that minimizes the absolute error between 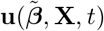 and ũ(**X**, *t*)

### 2.2 Constitutive and equilibrium equations

Equilibrium equations in *κ*_***β***_(*ℬ*) must be satisfied by the Cauchy stress **T**, which may be written, in absence of accelerations and body forces, as

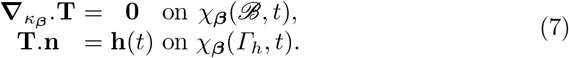

In Eq. 7, the equilibrium on the boundaries is only written where known tractions are applied as we will use these equilibrium equations to solve the identification problem and we do not want to introduce supplemental unknowns such as the reaction tractions on the boundaries where Dirichlet boundary conditions would be applied.

In this paper, we focus on compressible hyperelastic materials. It is assumed that their strain energy density function *Φ* can be written in an uncoupled form as

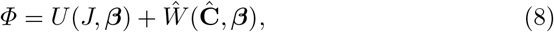

where *U* (*J*, ***β***) describes the volumetric response and *Ŵ* (**Ĉ**, ***β***) describes the deviatoric response.

The second Piola-Kirchhoff stress **S** can be written as

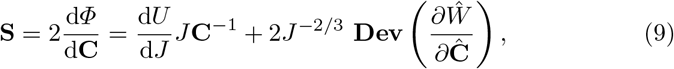

where

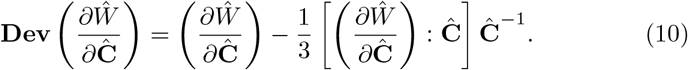

In the following, we introduce 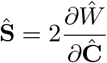 The Cauchy stress **T** can be written as

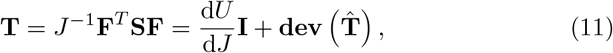

where 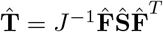, and

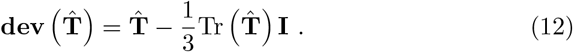

Moreover, we introduce

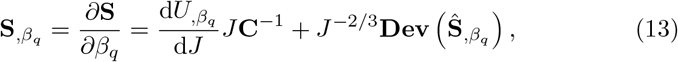

and

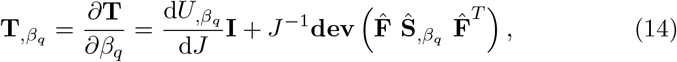

where

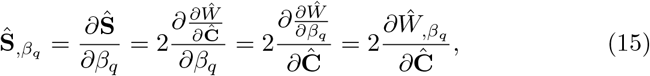

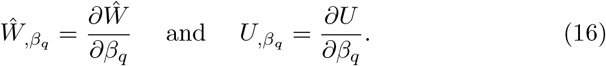

### 2.3 Definition of an intermediate configuration

#### 2.3.1 Decomposition of the deformation gradient

Let us start with a set of parameters ***β***^*o*^ which corresponds to an initial estimation of the unknown constitutive parameters based on existing literature on the same tissue for instance.

Unstressed “intermediate configuration” were traditionally used in the literature for the multiplicative split of the deformation gradient in finite strain plasticity [38]. We define here a stressed intermediate configuration *κ****β***^*o*^ (*B*) for which the solid with the vector of material properties ***β***^*o*^ is at equilibrium (Fig. 1). We emphasize that this is a different use of the “intermediate configuration” terminology than in finite strain plasticity [38].

**Fig. 1.**
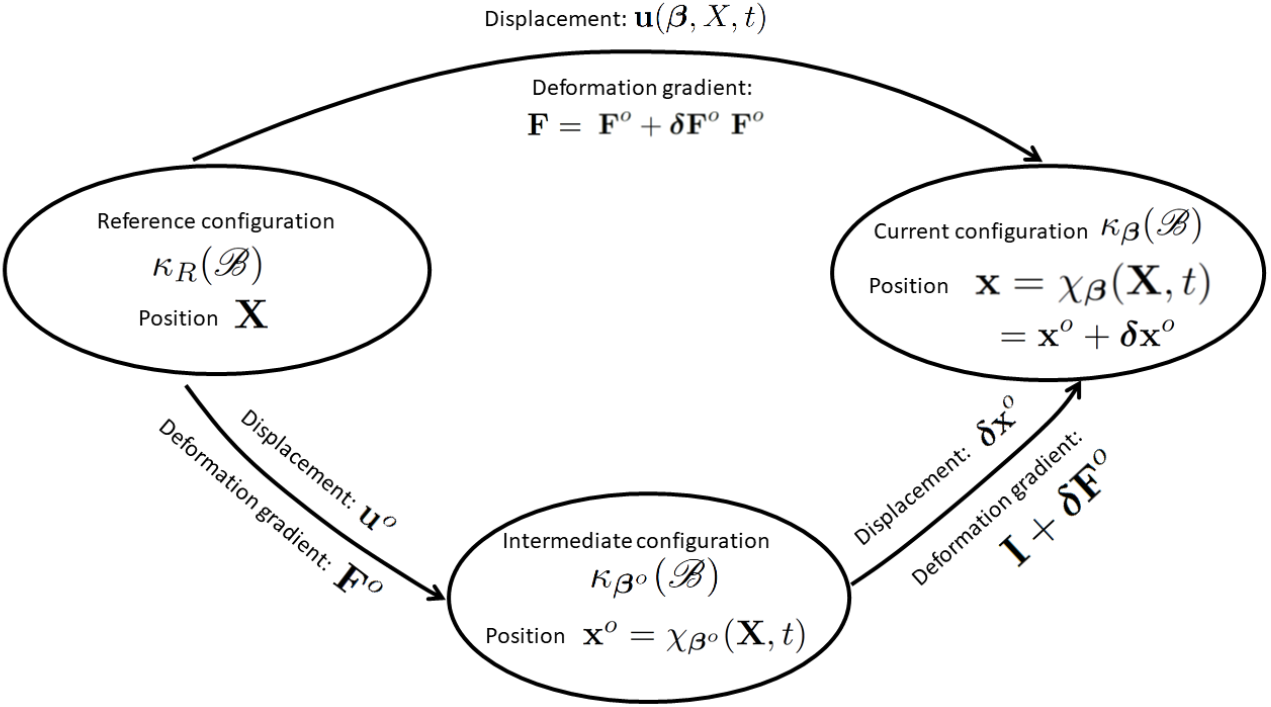
The relationship of the reference, intermediate and current configurations.

Then, let the position in the intermediate (stressed) configuration be denoted by **x**^*o*^ = *χ****β*** ^*o*^ (**X**, *t*), corresponding to displacement **u**^*o*^(**X**, *t*) = **x**^*o*^ −**X**. We assume that a small displacement ***δ*x**^*o*^ superimposed upon the large deformation **u**^*o*^, yields the current position **x** at time *t* for which the solid with a vector of material properties ***β*** = ***β***^*o*^ + ***δβ***^*o*^ is at equilibrium. We assume that ***δβ***^*o*^ is a small variation of ***β***^*o*^ such as

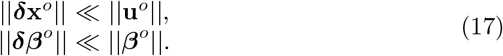

The small displacement ***δ*x**^*o*^ corresponds to the deformation between the intermediate configuration (at which the solid with ***β***^*o*^ material properties is at equilibrium) and the current configuration (at which the solid with ***β*** = ***β***^*o*^ + ***δβ***^*o*^ material properties is at equilibrium).

The current position can thus be written as

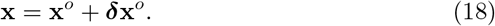

The deformation gradient associated with mappings from the reference to the intermediate is thus given by

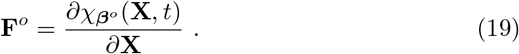

The deformation gradient representing a mapping from the intermediate configuration to the current configuration (corresponding to the variations of the configuration for a variation of material properties ***δβ***^*o*^) may be expressed as **I** + ***δ*F**^*o*^ where

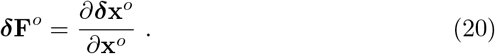

Then, gradients of the successive motions are obtained with the chain rule, such as

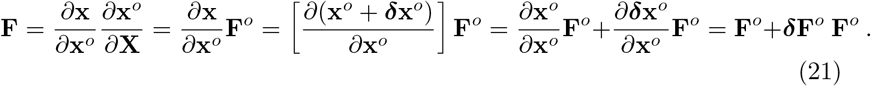

Then, the identification problem can be formulated as, find the values of ***δβ***^*o*^ that minimizes the error between ***δ*x**^*o*^ and **ũ** − **u**^*o*^.

#### 2.3.2 Cauchy stress tensor

Using Eq. 9 and Eq. 11 specifically with **F**^*o*^ deformation gradient, the second Piola-Kirchhoff stress **S**^*o*^ and Cauchy stress **T**^*o*^ in the intermediate configuration may be written respectively

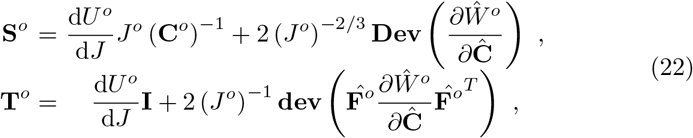

where *U*^*o*^ = *U* (*J*^*o*^, ***β***^*o*^) and 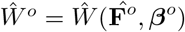.

Note that the Cauchy stress **T**^*o*^ must satisfy the equilibrium equations on the *κ****β***^*o*^ configuration (further denoted *κ*_*o*_) which may be rewritten

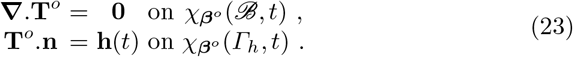

The second Piola-Kirchhoff stress **S** can be related to **S**^*o*^ using a Taylor expansion of first order. For that we substitute into Eq. 9, ***β*** = ***β***^*o*^ + ***δβ***^*o*^ and **C** = **C**^*o*^ + ***δ*C**^*o*^, which yields

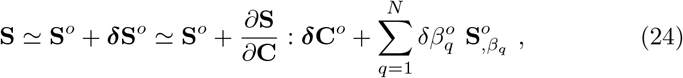

where

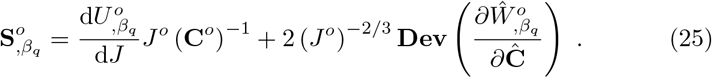

Neglecting the second order terms in the deformations from the intermediate to the current configuration, it may be written

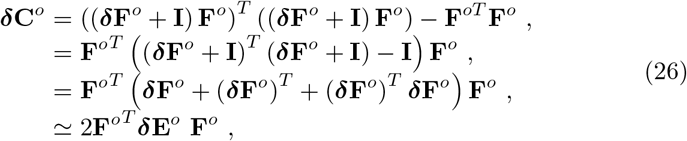

where 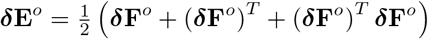 signifies the infinitesimal strain induced by a slight variation of material properties when ***δ*F**^*o*^ is small.

Finally,

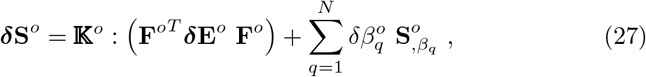

where,

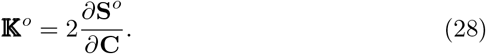

Next, we can derive the Cauchy stress **T** for the current configuration from **T**^*o*^ as

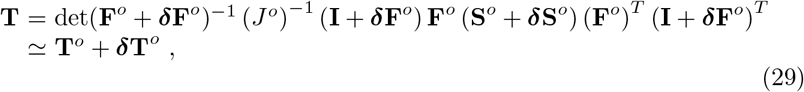

where

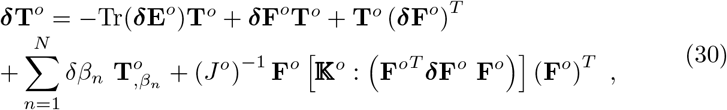

and

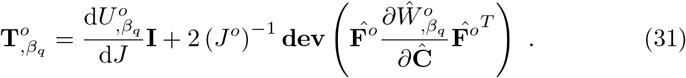

Equation 30 may be rewritten by introducing a 4^th^ order tensor 𝕃^*o*^ such as

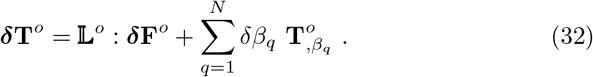

The components of the 𝕃^*o*^ tensor may be written such as

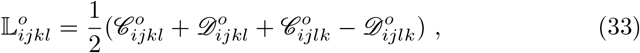

where, the summation convention being adopted for the repeated indices,

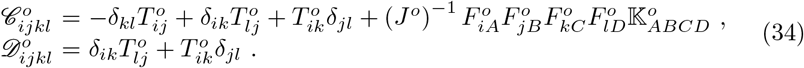

Equilibrium must be satisfied in the current configuration. Given that Eq. 23 is satisfied by **T**^*o*^ on the intermediate configuration, and assuming that the intermediate configuration is infinitesimally close to the current configuration (owing to ***δβ***^*o*^*/****β***^*o*^ ≪ 1), we can assume that the following equations are satisfied by ***δ*T**^*o*^,

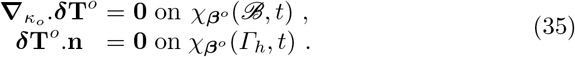

The equilibrium equations may be rewritten such as

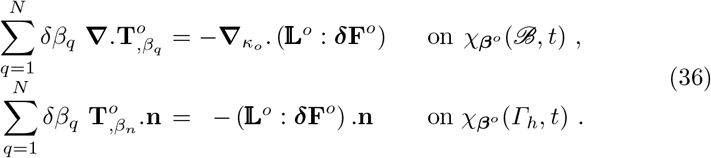

In the following, for the sake of simplification, the following abuse of notation will be used for the gradient and divergence in the *κ*_*o*_ configuration: 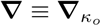.

### 2.4 The virtual fields method

***δ*T**^*o*^ should satisfy the equilibrium equations (Eq. 36) written just above. These equations can be written in their weak form such as

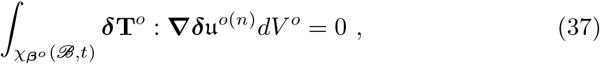

where ***δ***𝔲^*o*(*n*)^ is a virtual displacement field which equals zero on *χ****β***^*o*^ (*∂ℬ* \ *Γ*_*h*_) (boundary where traction are not applied). In ***δ***𝔲^*o*(*n*)^, the index (*n*) indicates that at least *N* virtual fields are necessary to establish a system of *N* equations of the *N* unknown material properties *β*_*n*_.

As full-field measurements are available, ***δ*F**^*o*^ and ***δ*E**^*o*^ from Eq. 30 can be replaced by their measures denoted respectively 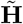 and 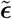, such as

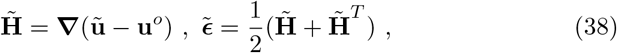

yielding

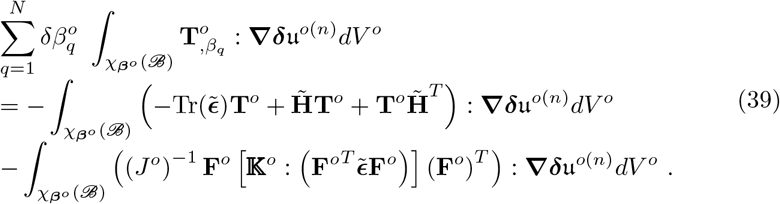

Equation 39 may be rewritten by introducing the 𝕃^*o*^ tensor such as

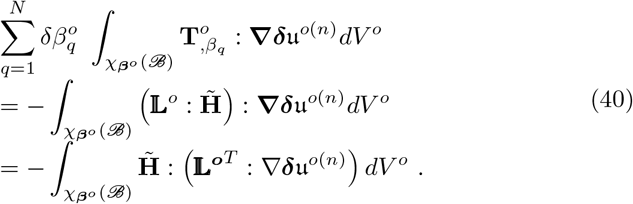

The previous equation can be written *N* times with *N* virtual fields ***δ***𝔲^*o*(*n*)^. The obtained system of equations may be written such as

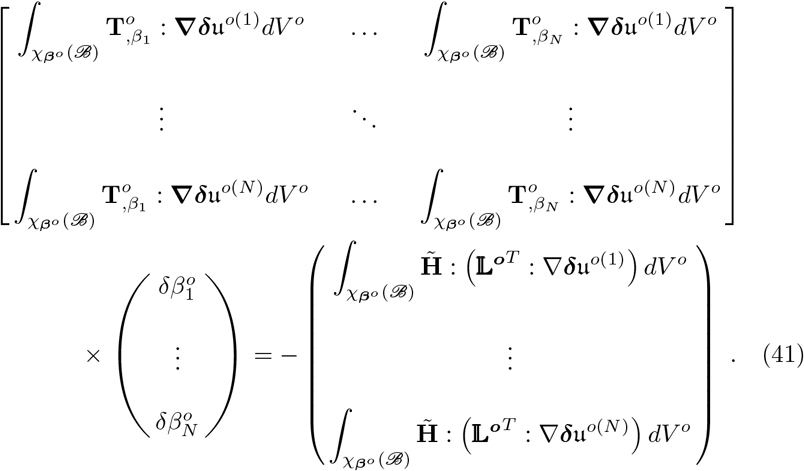

In the next section we provide a methodology to choose the set of *N* virtual fields.

### 2.5 Derivation of the virtual fields for parameter identification

In the previous subsection, we showed that the unknown constitutive parameters can be identified by solving a linear system of equations. To establish this system of equations, one needs to define *N* virtual fields denoted ***δ***𝔲^*o*(*n*)^, such that [**A**^*o*^] is invertible. The invertibility is ensured by relating each virtual field ***δ***𝔲^*o*(*n*)^ to the sensitivity of the Cauchy stress to each unknown parameter, denoted as 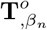 in Eq. 31 and introduced in Eq. 30, since it is assumed that the different 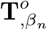 constitute a set of *N* linearly independent tensorial functions.

Therefore, a possible choice of *N* linearly independent virtual fields could simply be: 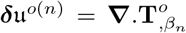. However, the virtual field must equal zero on *χ****β***^*o*^ (*∂ℬ*\*Γ*_*h*_) and remain continuous. To meet these requirements and benefit from the linear independence between the 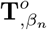 fields, we defined ***δ***𝔲^*o*(*n*)^ as the vectorial fields satisfying

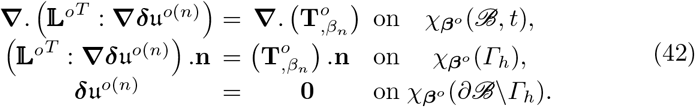

The virtual fields ***δ***𝔲^*o*(*n*)^ are eventually obtained by solving the linear elastic problems defined in Eq. 42 using the finite-element method (subsection 2.6). Let us introduce the *𝔏* linear operator such as 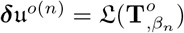 is the solution of Eq. 42.

Moreover using the integration by parts, we can write

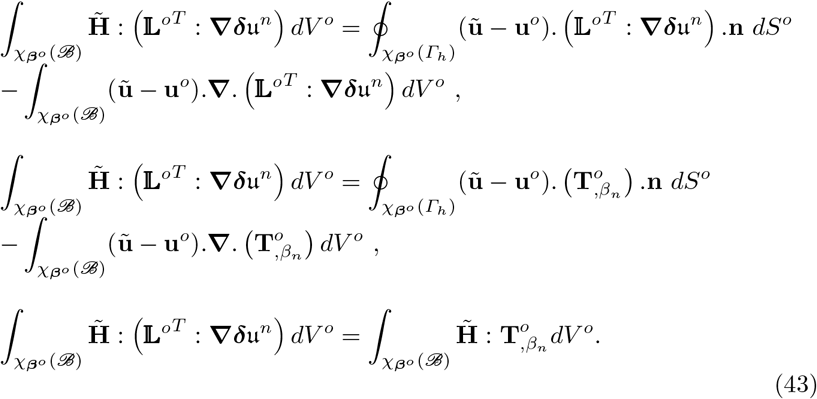

Then we can build the system of equations of Eq. 41 by replacing each ***δ***𝔲^*o*(*n*)^ by their expression coming from Eq. 42, and use also the simplifications derived in Eq. 43, yielding

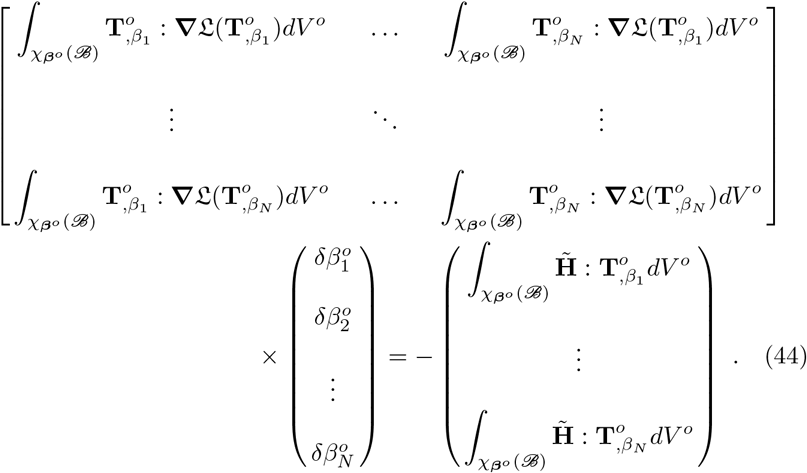

This system of equations can be rewritten

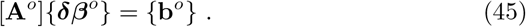

Then, the unknown constitutive parameters can be obtained as

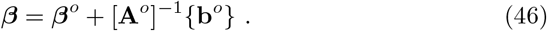

### 2.6 Finite-element implementation

The numerical implementation of the proposed method requires the use of finite-element analyses at two different levels:

1. find the intermediate configuration and the deformation gradient **F**^*o*^ by solving the partial differential equations (PDEs) of the forward problem (Eq. 23) with a set of parameters ***β***^*o*^,
2. compute integrals in Eq. 41 or Eq. 44 to identify the unknown constitutive parameters.

In this section, we propose a possible numerical implementation. As standard in finite-elements [39, 40, 41, 42]), the computational domain *ℬ* is discretised into a finite number of non-overlapping elements *e* ∈ 𝔼 such that

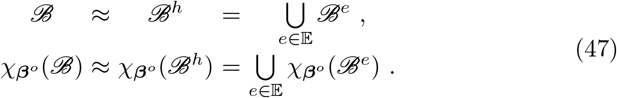

The fields are discretised using the following standard vectorial shape functions ***𝒩*** (**X**) as

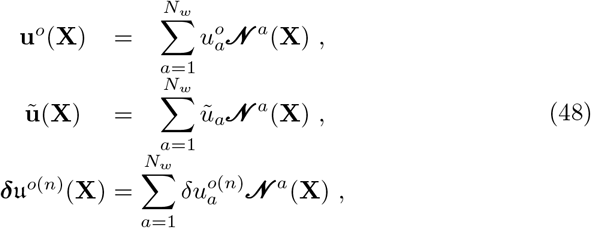

where 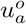 are nodal variables for the **u**^*o*^ field, *ũ*_*a*_ are nodal variables for the **u**^*o*^ field and 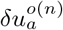 are nodal variables for the ***δ***𝔲^*o*(*n*)^ field.

A standard finite-element discretisation enables the following discrete form for the components of Eq. 44,

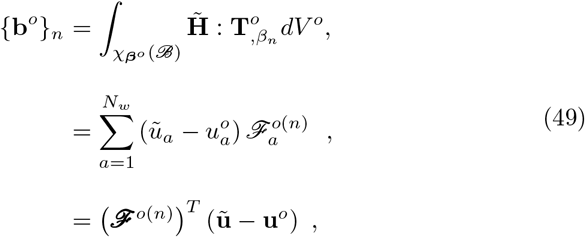

where ***ℱ*** ^*o*(*n*)^ is obtained such as

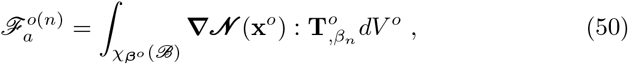

and

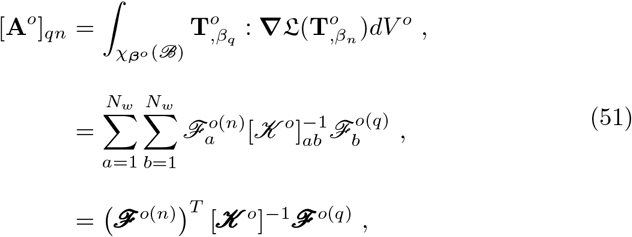

where ***𝒦*** ^*o*^, which is the stiffness matrix needed to solve the elastic problem of Eq. 42 to derive the virtual fields, and which satisfies ***𝒦*** ^*o*^***δ***𝔲^*o*(*n*)^ = ***ℱ*** ^*o*(*n*)^, is obtained such as

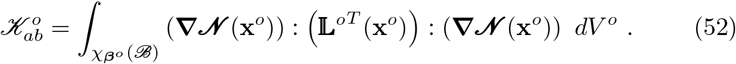

Finally, the expression of [**A**^*o*^] and {**b**^*o*^} with this finite-element discretisation are

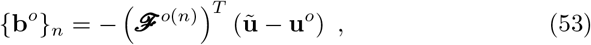

and

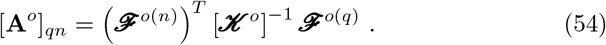

### 2.7 Convergence of parameter identification

Based on the details discussed in the previous subsection, it is possible to derive ***δβ***^*o*^ from the choice of an initial guess of ***β***^*o*^ by solving Eq. 45. The question that remains to be solved is how to choose ***β***^*o*^. In practice, an initial estimation of the unknown constitutive parameters based on existing literature on the same tissue can be used to initiate the resolution. However, this does not guarantee the criterion in Eq. 17 is satisfied. Then, a non infinitesimal deviation between the intermediate configuration and the reference configuration would cause Eq. 36 to not be satisfied on *χ****β***^*o*^ (*ℬ, t*). Therefore, the concept of “intermediate configuration” has to be iterative. From the choice of a first set of parameters ***β***^*o*^, an intermediate configuration can be found by solving the forward problem in Eq. 23. Evaluating [**A**^*o*^] and {**b**^*o*^} (from equations 53 and 54) using the obtained 𝕃^*o*^ and 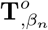 expressions (from equations 33 and 31) provides an update for ***β***^*o*^. The process is repeated until the deviation between **u**^*o*^ and **ũ** becomes small enough (Fig. 2).

**Fig. 2.**
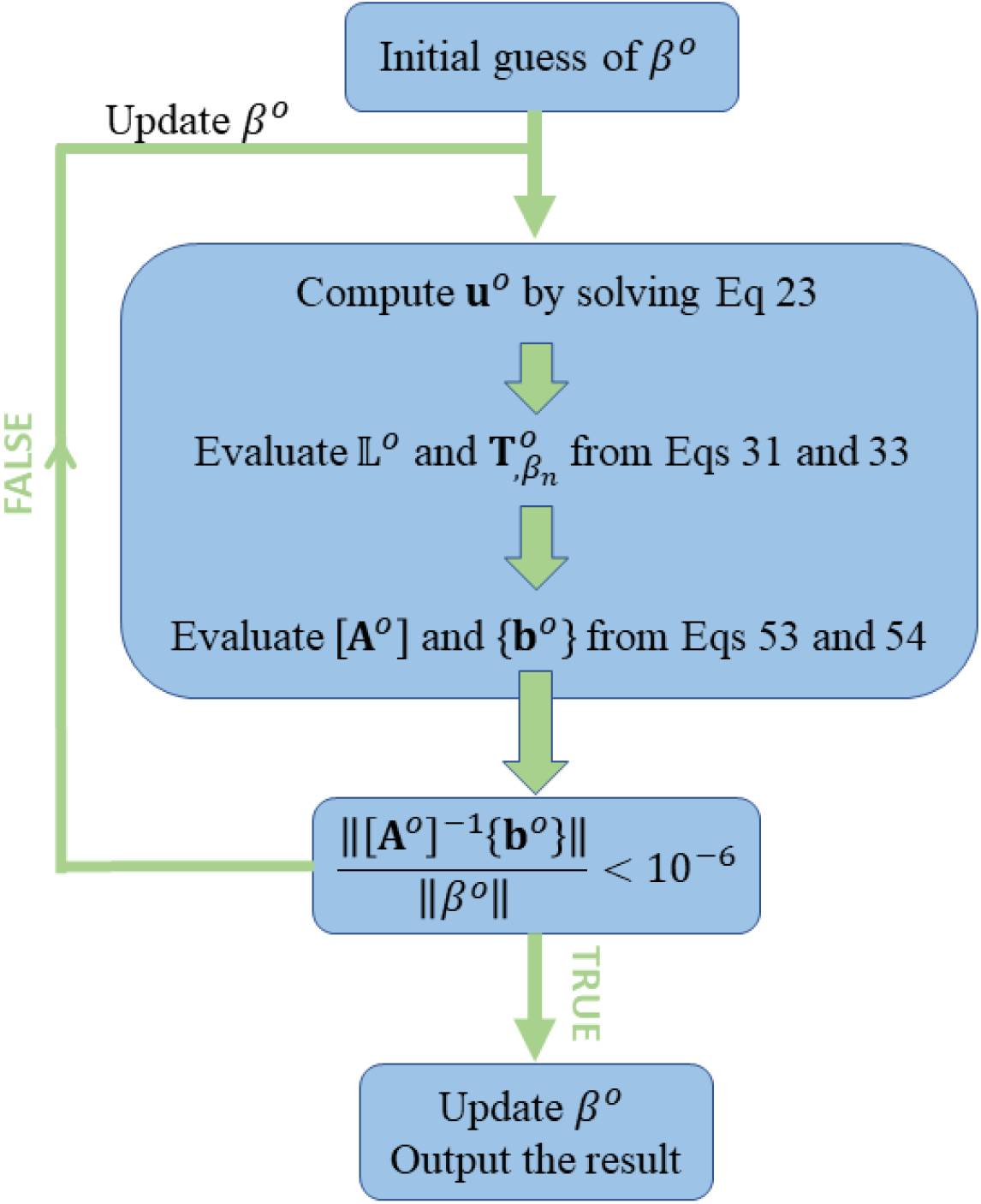
Flowchart of the proposed inverse method.

The convergence criterion of the inverse algorithm is that the relative difference between the current estimation of material properties ***β***^*o*^ and its update [**A**^*o*^]^*−*1^{**b**^*o*^} is less than the tolerance delta

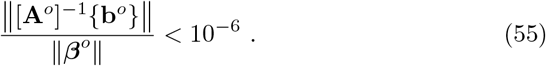

Although deriving analytically the convergence rate is not possible, we verified numerically in the following results that a quadratic convergence was obtained for cases where existence and uniqueness of the solution are guaranteed.

### 2.8 Examples of hyperelastic constitutive models

In the following sections, we applied the VFM method and algorithm in Fig. 2 to determine the parameters of 3 hyperelastic strain energy potentials commonly used to describe collagenous tissues.

#### 2.8.1 Neo-Hookean model

For the Neo-Hookean constitutive model, the strain energy density function is

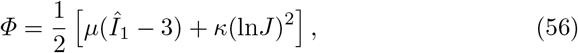

where *Î*_1_ = Tr **(Ĉ**)= *J* ^*−*2*/*3^Tr (**C**).

The strain energy density function depends linearly on two unknown constitutive parameters denoted *µ* and *κ*. Then in an identification problem, the vector of unknown parameters would be: *β* = (*µ, κ*).

The second Piola-Kirchhoff stress is written

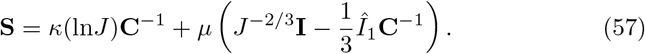

The associated Cauchy stress is written as

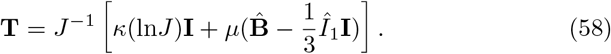

The sensitivity of the second Piola-Kirchhoff stress to each parameter is written

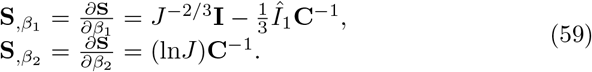

The sensitivity of the Cauchy stress to each parameter is written

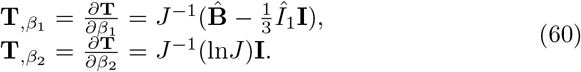

Therefore,

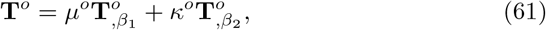

with

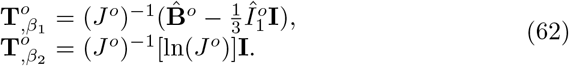

Moreover, we have

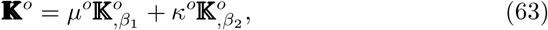

with

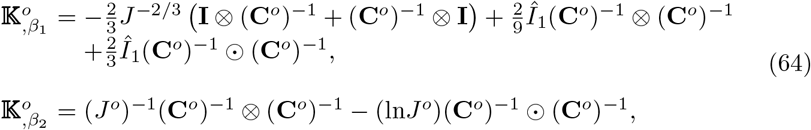

where 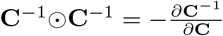 and ⊗ represents the dyadic multiplication symbol.

#### 2.8.2 Mooney-Rivlin model

For the Mooney-Rivlin constitutive model, the strain energy density function is

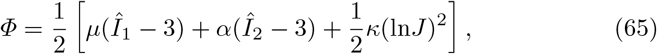

where *Î* _2_ = *J* ^*−*4*/*3^*I*_2_ = *J* ^*−*4*/*3^Tr (**Ĉ)** = *J* ^*−*2*/*3^Tr (**C**).

The strain energy density function depends linearly on three unknown constitutive parameters denoted *µ, α* and *κ*. Then in an identification problem, the vector of unknown parameters would be: *β* = (*µ, α, κ*).

The second Piola-Kirchhoff stress is written

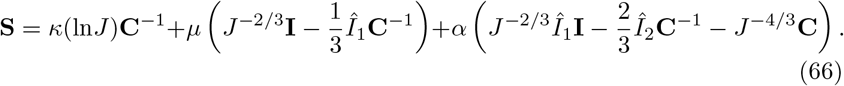

The associated Cauchy stress is written

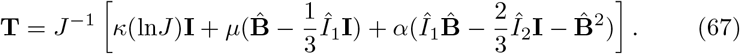

The sensitivity of the second Piola-Kirchhoff stress to each parameter is written

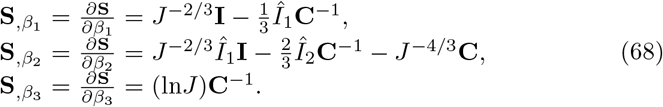

The sensitivity of the Cauchy stress to each parameter is written

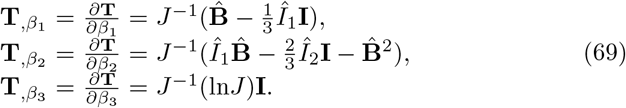

Therefore,

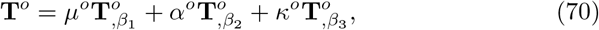

with

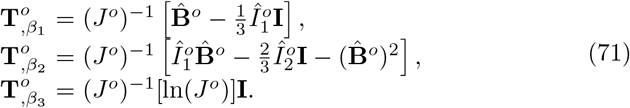

Moreover, we have

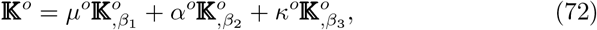

with

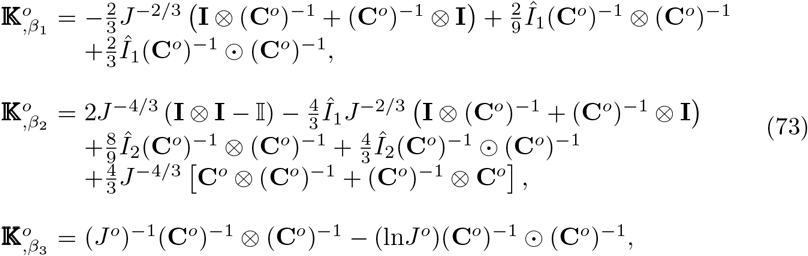

where 𝕀 is the fourth order identity tensor.

#### 2.8.3 Veronda-Westmann model

In the two previous constitutive models, the strain energy density function depends linearly on the material properties. In this subsection, the Veronda-Westmann model [43] which involves an exponential function of a material property introduced. The strain energy density function is written such as

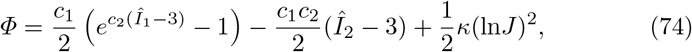

where *c*_1_, *c*_2_ and *κ* are material properties. Note that *c*_2_ is a parameter controlling the nonlinear behavior of the Veronda-Westmann solid, and *µ* = *c*_1_*c*_2_ is the shear modulus controlling the linear behavior of the Veronda-Westmann solid. In the following, we rewrite the model such as

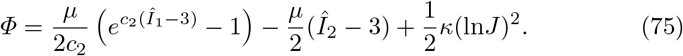

The strain energy density function depends linearly on two unknown consti-tutive parameters denoted *µ* and *κ*, and non linearly on the unknown parameter *c*_2_. In the identification problem, the vector of unknown parameters would be written: *β* = (*µ, c*_2_, *κ*).

The second Piola-Kirchhoff stress is written

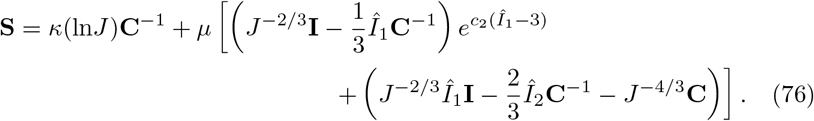

The associated Cauchy stress is written

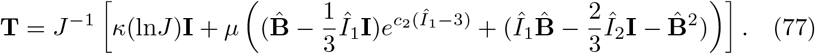

The sensitivity of the second Piola-Kirchhoff stress to each parameter is written

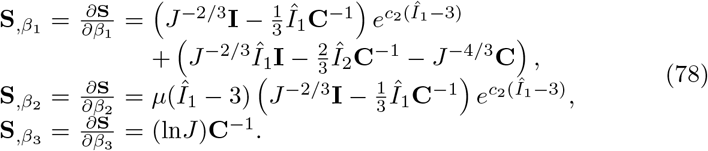

The sensitivity of the Cauchy stress to each parameter is written

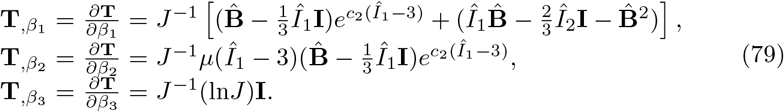

Then we approximate **T**^*o*^ such as

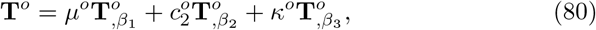

with

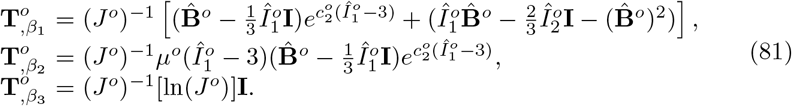

Moreover, we also approximate

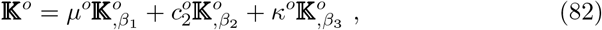

with

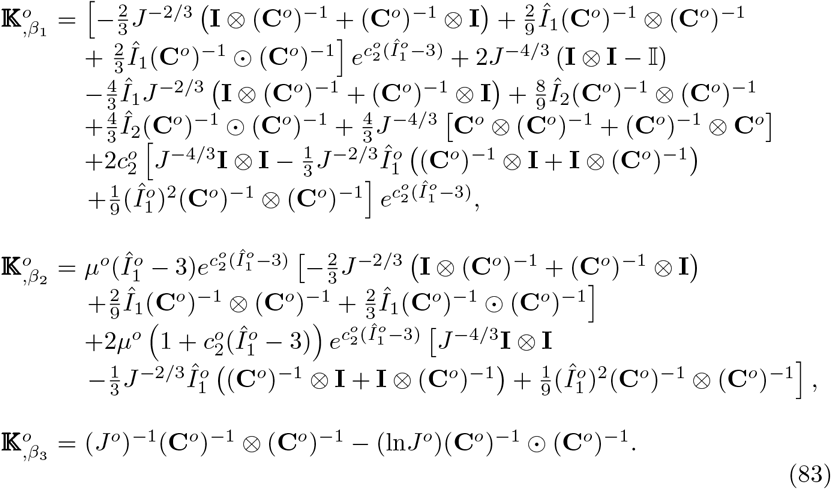

### 2.9 Case study for verification

Using the previous formulas derived for the Neo-Hookean, the Mooney-Rivlin and the Veronda-Westmann models, it is straightforward to implement the novel VFM method and apply it to solve identification problems based on full-field data. Note also that the approach also extends naturally to more complex strain energy density functions involving exponential functions with invariants *I*_4_ and *I*_6_ [28] for instance. In the next section we show results for the 3 constitutive models, which were previously introduced, and discuss the convergence and the feasibility of the proposed approach in the case of compression and tension tests onto 2 simple geometries (Fig. 3):

**Fig. 3.**
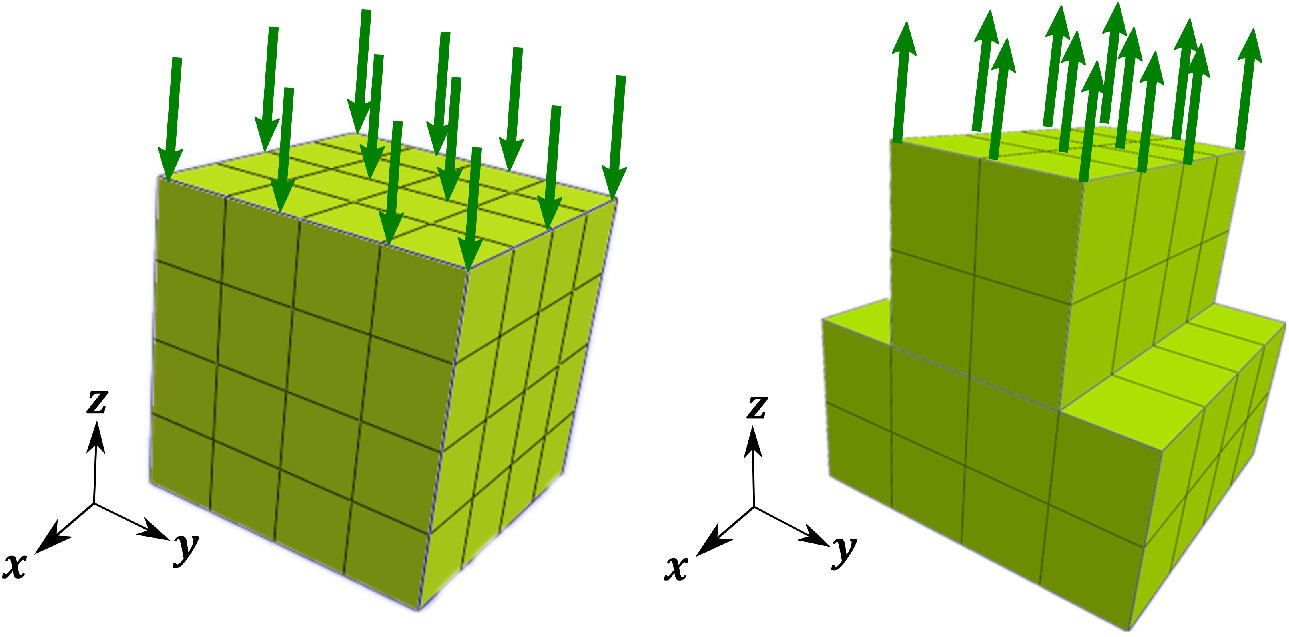
The geometry and finite-element meshes used to verify the implementation of the novel VFM approach.

1. Compression is applied on a cube of 1 cm edge which is fully fully fixed on its bottom face (Fig. 3a). For the sake of verification of the approach, the cube is simply discretized uniformly by 4× 4 ×4 elements.
2. We also consider another geometric model as presented in (Fig. 3b) where we apply tension on the top face and fix the bottom face.

In both cases, full-field displacement measurements were simulated by solving finite-element models using the open source software FEBio [44]. For each case, the full-field displacement measurements were simulated with a set of target material properties, named the target, and the proposed VFM approach was employed to recover the target for the Neo-Hookean, the Mooney-Rivlin and the Veronda-Westmann model, using different initialization values to assess the convergence of the method.

### 2.10 Case study for validation

The proposed VFM method will be used in this case study to determine the material properties of the LC, a connective tissue structure in the ONH of great interest to researchers studying development and progression of glaucoma [45]. The LC in humans is approximately 1.5 mm - 2.0 mm in diameter and 450 microns thick and is located in the back of the eye. The mechanics of the LC is thought [46] to play an important role in mediating the progressive vision loss associated with glaucoma, a degenerative disease that is a leading cause of blindness worldwide [47]. As shown in Figure 4, the LC is situated beneath the prelaminar neural tissue (PLNT). We further split the LC into an anterior region (ALC), 250 *µ*m posterior to the anterior posterior surface, and a posterior region (PLC), which includes the remainder of the imageable volume of the LC.

**Fig. 4.**
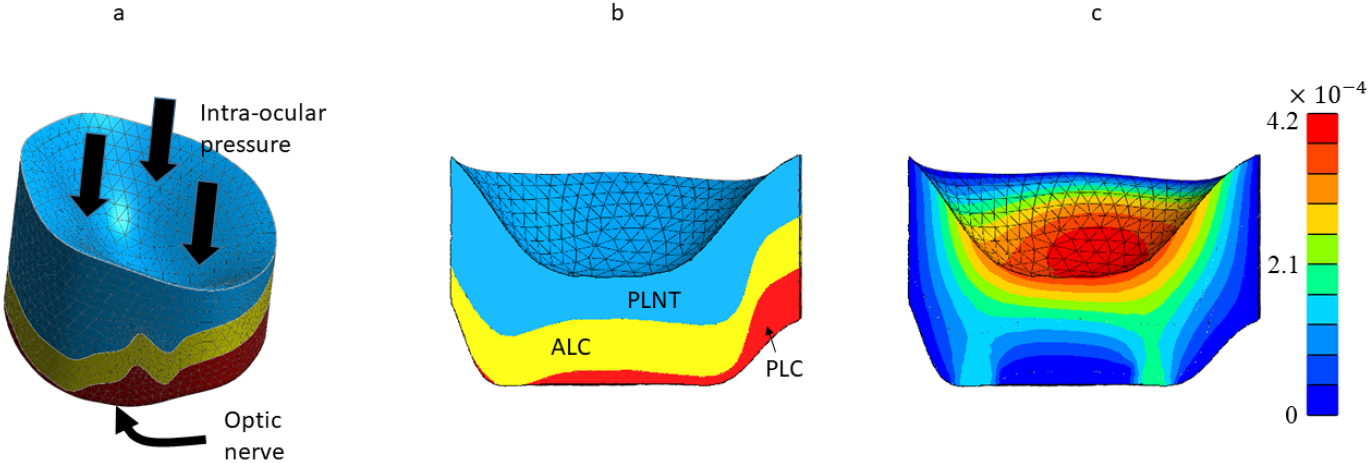
(a) Schematic showing the solid model of the optical nerve head (ONH). The anterior (top) face is subjected to the intra-ocular pressure. The posterior (bottom) face is in direct contact with the optic nerve. (b) Cross sectional view of the ONH solid model showing the 3 regions with different material properties: the prelaminar neural tissue (PLNT), the anterior lamina cribosa (ALC) and the posterior lamina cribosa (PLC). (c) Displacement field caused by reducing the intraocular pressure of 10 mmHg (colorbar in mm).

Spectral domain OCT imaging (Heildelberg Engineering) was applied to acquire 24 radial scans centered about the ONH of the left eye of a glaucoma patient. The OCT imaging was performed at Johns Hopkins University’s Wilmer Eye Institute in the Glaucoma Center of Excellence, and was approved by the appropriate Institutional Review Board. The following structural features were marked in the 24 radial scans to segment the tissue structures of the ONH as described in Midgett et al. [48]: Bruch’s membrane opening, the anterior boundary of the PLNT, the anterior LC surface, and the boundary of the imageable volume below the anterior LC surface. The manual marking were imported into Cubit (Coreform, Orem, UT, USA) to construct surface geometries using closed splines. The PLNT, ALC, and PLC volumes were defined by extruding a cylinder from Bruch’s membrane posteriorly to intersect the anterior PLNT surface and the anterior LC surface (for the PLNT), the anterior ALC surface and a surface positioned 250 microns posterior to the anterior ALC surface (for the ALC), and that posterior surface and a surface marking the end of the imageable volume of the LC (see Figure 4a). The final solid volume was meshed in Cubit linear 4-node tetrahedral elements and exported into FEBio, where the linear elements were converted to 10-node quadratic tetrahedral elements [49] to avoid mesh locking and improve accuracy. The compressible Neo-Hookean constitutive model (Eq. 56) was chosen for all three materials.

To validate the present VFM method, we simulated a displacement field by solving the forward problem with target parameter values, which are reported in Tab. 2. In the forward problem, zero displacement boundary conditions were applied to the posterior and lateral surfaces. To account for the 10 mmHg pressure decrease, a pressure boundary condition was applied to the ONH surface, with a magnitude of −10 mmHg. The induced displacement is shown in Figure 4c.

Eventually the proposed VFM approach was employed to process the simulated displacement fields and recover the target compressibility modulus values for each separate material.

## 3 Results

### 3.1 Neo-Hookean model

In this section, we tested the feasibility of the proposed VFM approach for the compressible neo-Hookean model using the 10-mm-edge cubic specimen under compression shown in Fig. 3a. The cube was discretized uniformly by 4 × 4 × 4 elements. We applied a deformation of 2 mm to the cubic specimen. The target values of material properties were: Young’s modulus *Ē* = 10Mpa and Poisson’s ratio 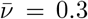, which corresponds to 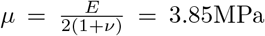 and 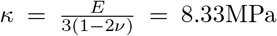 We choose three different initial guess pairs (*E*^*o*^ = 40MPa, *ν*^*o*^ = 0.45), (*E*^*o*^ = 5MPa, *ν*^*o*^ = 0.45) and (*E*^*o*^ = 40MPa, *ν*^*o*^ = 0.15). The convergence plots for each case are shown in Fig. 5 for the objective function value and in Fig. 6 for the evolution of the obtained parameter values. The material parameters were recovered regardless of the initial guesses. Convergence was reached after 4 to 6 iterations indicating the quadratic convergence of the new VFM approach.

**Fig. 5.**
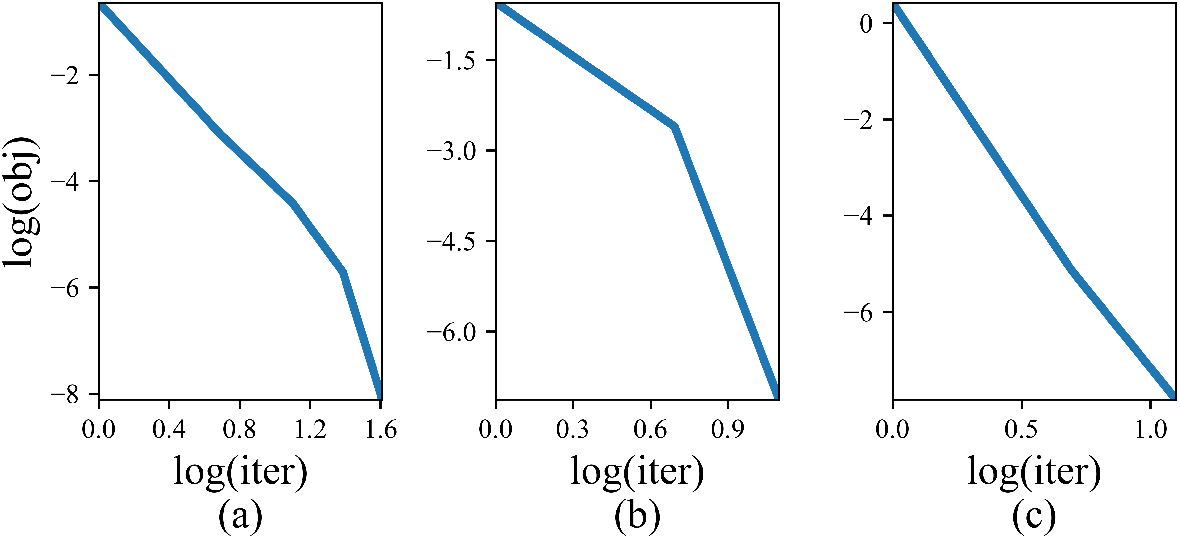
Evolution of the objective function relative value ‖**u**^*o*^ −**ũ**‖ /‖ **ũ** ‖ (log-log representation) for the identification of each unknown parameter of the neo-Hookean model with the novel VFM approach, using initial guesses *E*^*o*^ = 40MPa, *ν*^*o*^ = 0.45 in (a), *E*^*o*^ = 5MPa, *ν*^*o*^ = 0.45, in (b) and *E*^*o*^ = 40MPa, *ν*^*o*^ = 0.15 in (c).

**Fig. 6.**
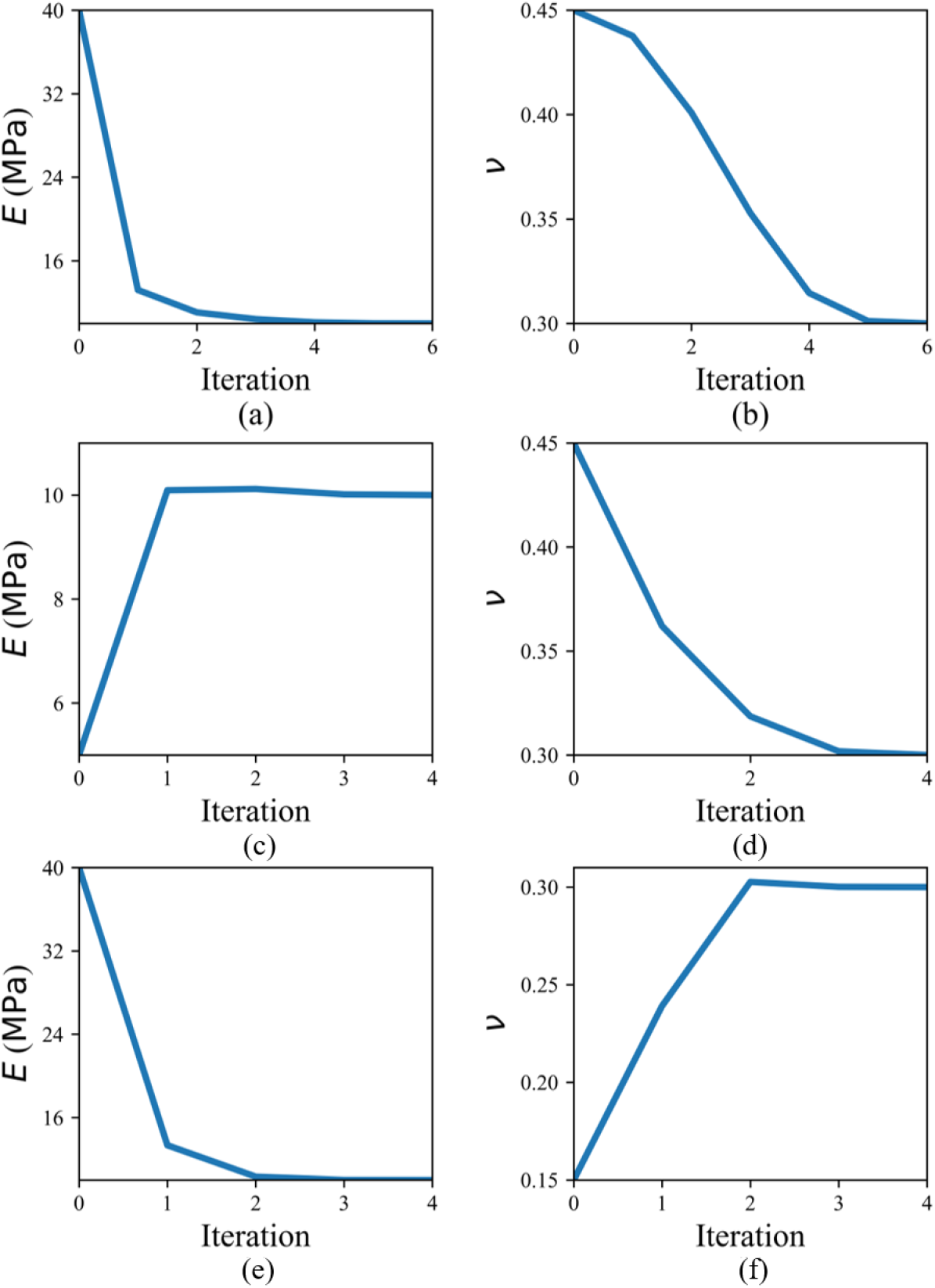
Convergence plots for the identification of each unknown parameter of the Neo-Hookean model with the novel VFM approach, using initial guesses *E*^*o*^ = 40MPa, *ν*^*o*^ = 0.45 in (a) and (b), *E*^*o*^ = 5MPa, *ν*^*o*^ = 0.45 in (c) and (d) and *E*^*o*^ = 40MPa, *ν*^*o*^ = 0.15 in (e) and (f).

### 3.2 Mooney-Rivlin model

Then, we tested the feasibility of the proposed VFM approach for the Mooney-Rivlin model, which includes three unknown material parameters. In this case, the material property vector was denoted *β* = [*µ, α, κ*]. Note that when *α* = 0, the Mooney-Rivlin model simplifies into the neo-Hookean model. For the verification of the VFM, we used again the cubic model under compression. The target material properties were set to 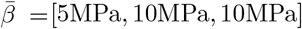. We chose two different sets of initial guess to evaluate the material identification problem. Convergence plots for each material parameters are shown in Fig. 7. The material parameters were recovered regardless of the initial guesses. Convergence was reached after 4 to 6 iterations indicating indicating the quadratic convergence of the new VFM approach with the Mooney-Rivlin model.

**Fig. 7.**
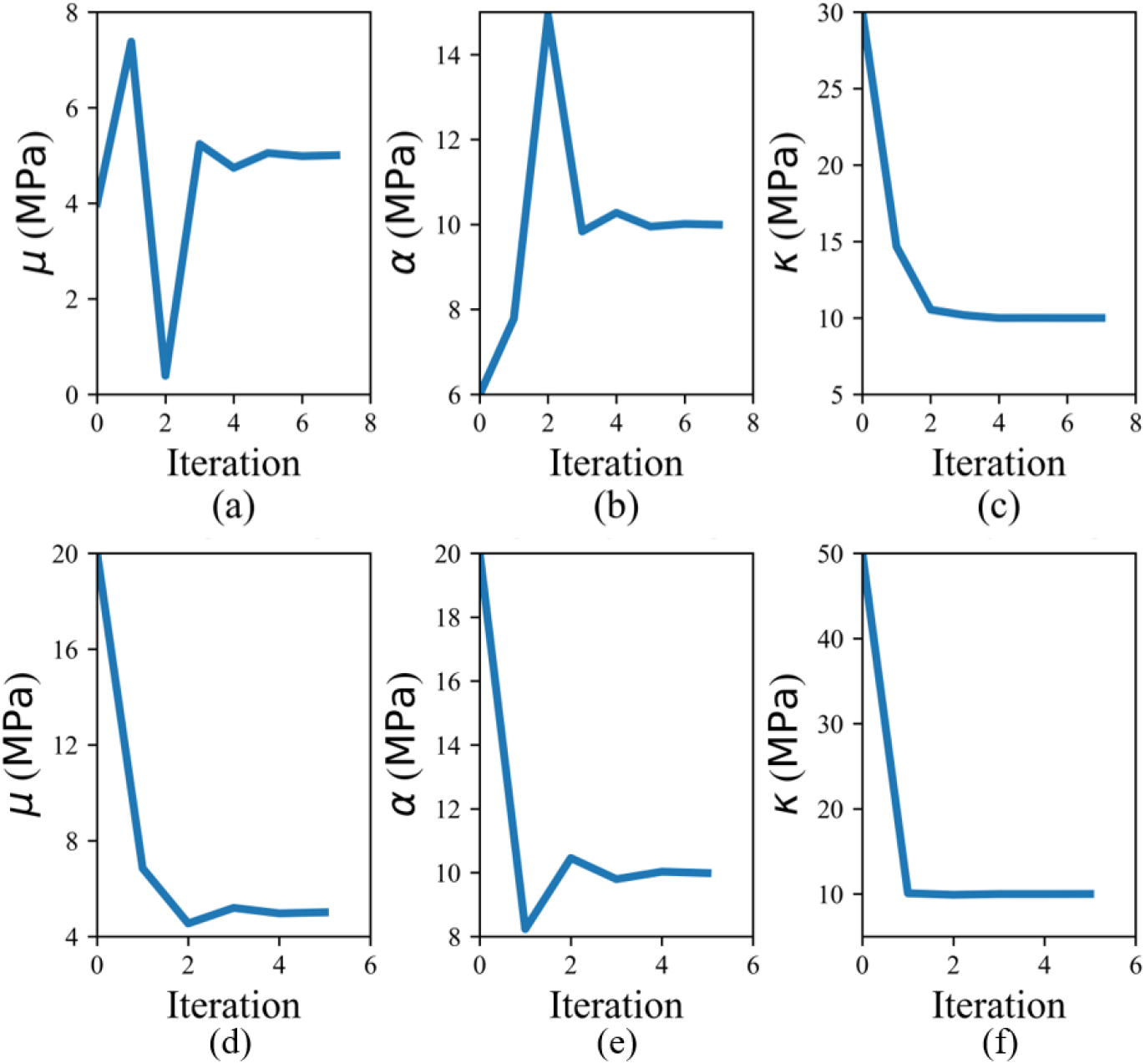
Convergence plots of the novel VFM approach for each parameter of the Mooney-Rivlin model using two sets of initial guesses: (a, b and c): *β*^*o*^ = [4MPa, 6MPa, 30MPa] and (d, e and f) *β*^*o*^ = [20MPa, 20MPa, 50MPa].

### 3.3 Veronda-Westmann model

Usually, to estimate the material parameters of the Veronda-Westmann model, displacement data with multiple loading stages are required due to the exponential term. In this work, we attempted to use only one loading stage to estimate the material parameters in the Veronda-Westmann model using the novel VFM approach. For that, we used the geometric model on the right hand side of Fig. 3. We applied tension on the top face and fix the bottom face. A deformation of 2 mm was applied. The target material parameters were 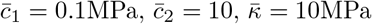. We used two different initial guesses. The initial guesses of the material parameters were 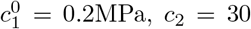, *c*_2_ = 30, *κ* = 20MPa. Another initial guess is set to 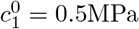, *c*_2_ = 5, *κ* = 5MPa

The convergence performance of each material parameters is shown in Fig. 8. We observed that 12 iterations were needed this time to reach convergence. Although the proposed VFM approach was capable of estimating material properties of the Veronda-Westmann solid using only one loading stage with a quadratic convergence, the convergence rate was slower due to the nonlinearity of the constitutive equations.

**Fig. 8.**
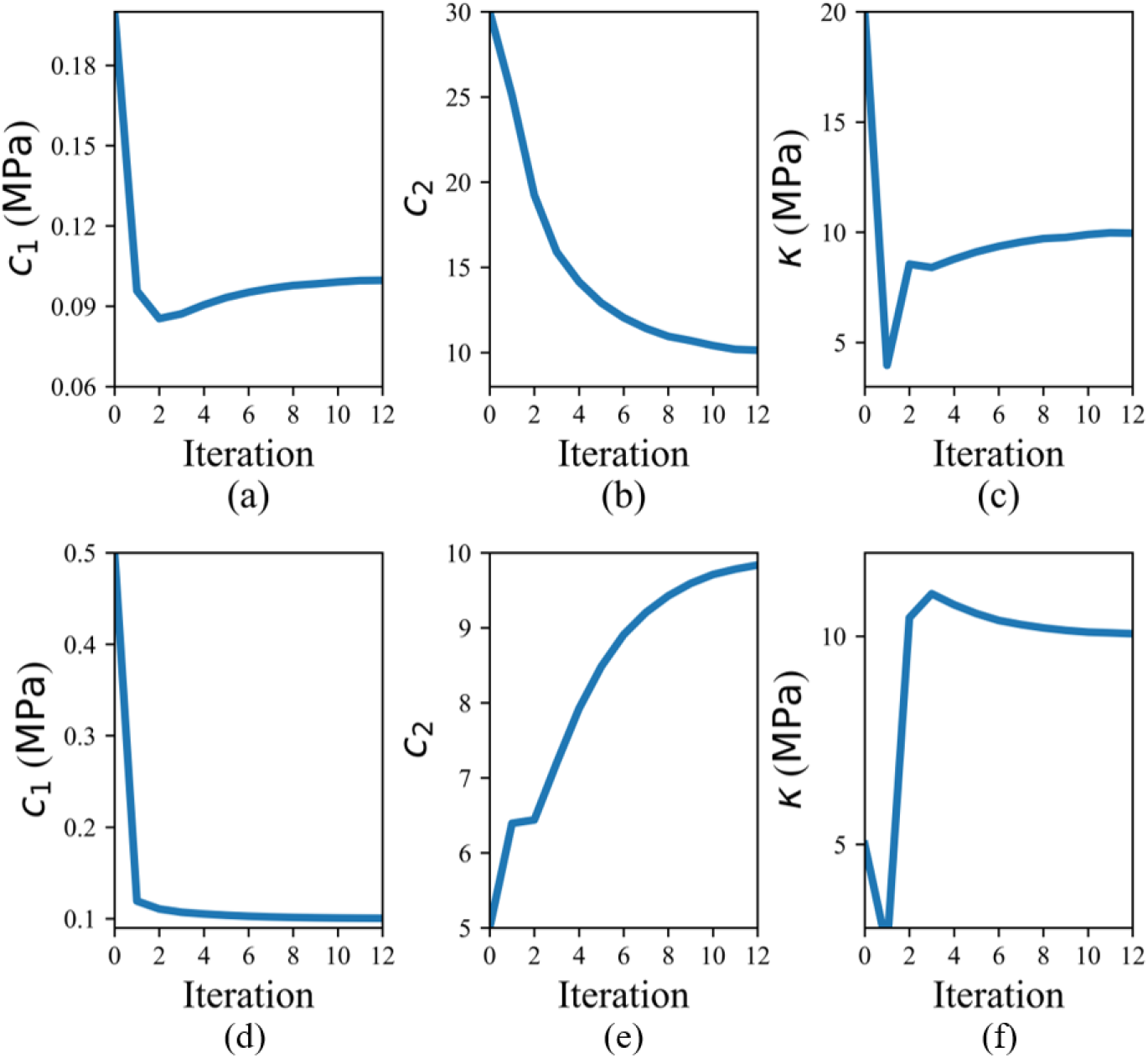
Convergence plots of the novel VFM approach for each parameter of the Veronda-Westmann model with initial guesses 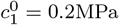, *c*_2_ = 30, *κ* = 20MPa in (a), (b) and (c), and initial guesses 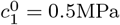, *c*_2_ = 5, *κ* = 5MPa in (d), (e) and (d).

### 3.5 Sensitivity to the compressibility parameters

Most biological tissues are nearly or truly incompressible. Thus, the sensitivity of the proposed method to the compressibility should be investigated. For incompressible materials, fewer material properties have to be identified compared to the associated compressible constitutive model. Thus, we considered the Neo-Hookean model as an example and focused on compressible cases with Poisson’s ratio *ν* = 0.45 and Young’s modulus *Ē* = 10MPa, which corresponds to 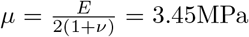 and 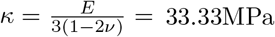.

We chose two different initial guess pairs (*E*^*o*^ = 5MPa, *ν*^*o*^ = 0.499), (*E*^*o*^ = 20MPa and *ν*^*o*^ = 0.499). The convergence plots for each case are shown in Fig. 9. We observed that it took 8 iterations for convergence, slightly more than the more compressible cases where *ν* was taken equal to 0.3.

**Fig. 9.**
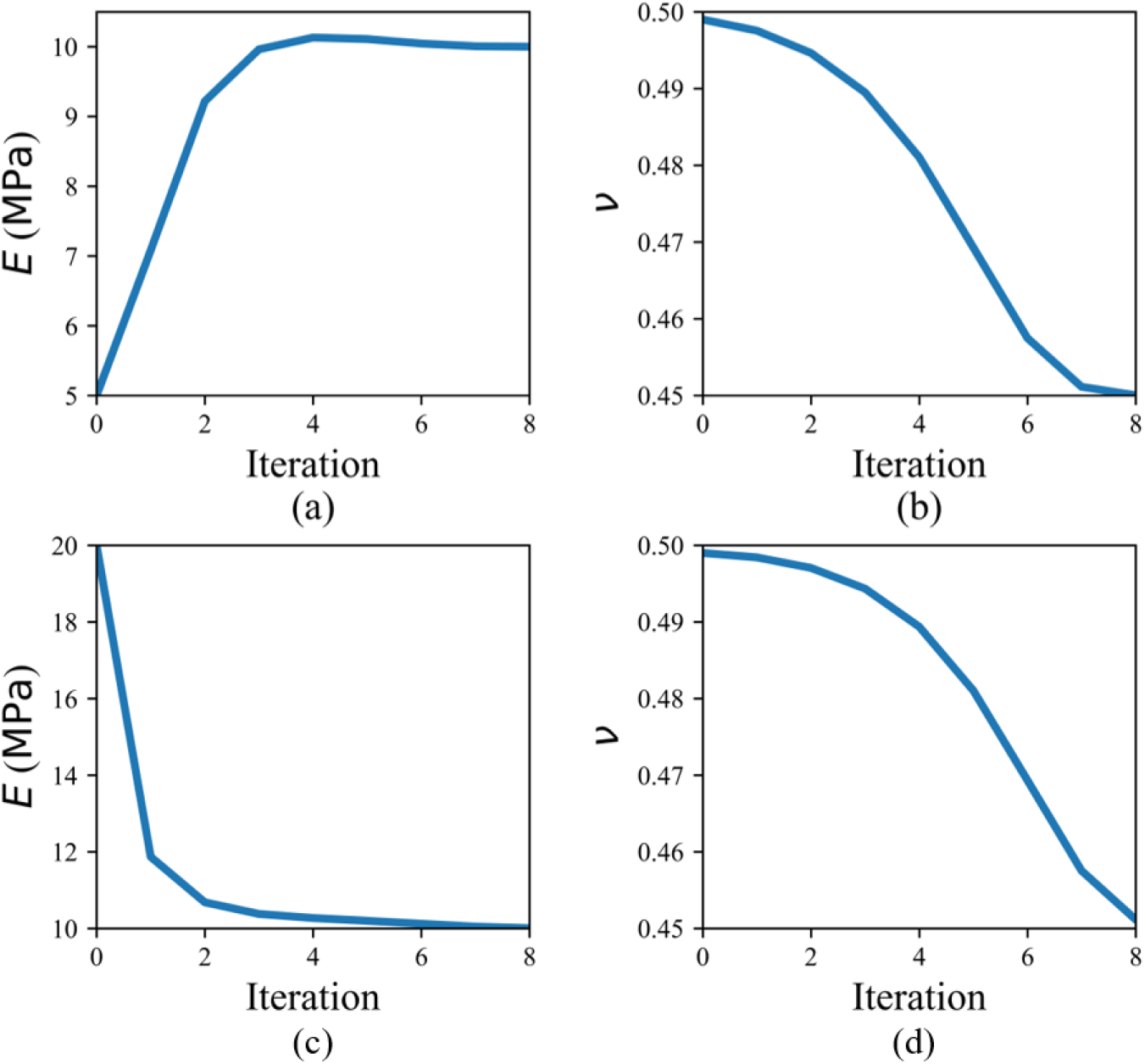
Convergence plots for the identification of each unknown parameter of the neo-Hookean model with Poisson’s ratio *ν* = 0.45, using initial guesses *E*^*o*^ = 5MPa, *ν*^*o*^ = 0.499 in (a) and (b), and *E*^*o*^ = 20MPa, *ν*^*o*^ = 0.499 in (c) and (d).

### 3.5 Case study for validation

The patient-specific geometry of the ONH, which was reconstructed using OCT data (Figure 4), was split into 3 parts with different compressibility moduli. It was assumed that each part had the same shear modulus *µ* = 0.2 MPa. Identifying the 3 unknown compressibility moduli with the VFM requires to define 3 independent virtual fields across the volume of interest. As any kinematically admissible virtual displacement field is a possible option, there are an infinite number of choices for these virtual fields. However, only very specific virtual fields ensure that the final system of equations is well conditioned and it may be cumbersome to determine these virtual fields by trial and errors. Here we applied the new VFM approach, where systems of equations are automatically built, without trial and error verifications, on the patient-specific geometry of the lamina cribrosa. As our VFM approach is iterative, values need to be chosen for initializing the compressibility moduli at the first iteration. We drew randomly these initial values within the range [1–5] MPa. The VFM always converged towards the target moduli. The largest error was obtained when the initial values were 5 MPa for the 3 compressibility moduli. Results obtained in that case are reported in Tab. 2. Errors below 1% can be reached after only 4 iterations of the new VFM approach.

## 4 Discussion

In this paper, we proposed a novel general framework to solve identification problems in hyperelasticity with the VFM. For that, we employed the finite-element method and generated the virtual fields by solving novel equations derived in this paper. The main advantage of the new approach for generating the virtual fields is that it can be applied to any kind of geometric shapes.

**Table 2.** Results obtained with the novel VFM for identifying compressibility modulus values of the 3 separate regions in the ONH using simulated data.

| | Reference $\kappa$ | Identified $\kappa$<br>after 4 iterations | relative<br>error | Identified $\kappa$<br>after 5 iterations | relative<br>error |
| --- | --- | --- | --- | --- | --- |
| PLNT | 2.29 MPa | 2.31 MPa | 0.82% | 2.284 MPa | 0.35% |
| ALC | 2.5 MPa | 2.51 MPa | 0.55% | 2.495 MPa | 0.22% |
| PLC | 2.71 MPa | 2.72 MPa | 0.37% | 2.706 MPa | 0.08% |

The VFM was previously applied to identify multiaxial hyperelastic properties of murine dissecting aneurysm samples [29, 30, 25] or hyperelastic properties in human ocular tissues [23]. This permitted significant progress especially in the characterization of heterogeneous material properties of lesions exhibiting complex morphologies, with different regions characterized by localized changes in tissue composition, microstructure, and properties. This was rendered possible by combining extension-distension data with full-field multimodality measurements of wall strain and thickness to inform the inverse material characterization using the virtual fields method.

Latest key advances were the use of a digital volume correlation data [23, 25] that allowed for characterization of properties in the bulk of soft tissues. However, despite these progresses, generic rules had never been defined for the choice of virtual fields. For instance, in [23], analytical expressions of virtual fields were written in the cylindrical coordinate system using the *arctan* function. In [29], virtual fields were also defined analytically, permitting to yield expressions generalizing the Laplace’s law for pressurized membranes.

The main difficulty of these previous studies was to ensure that the choice of virtual fields can provide enough equations to identify all the unknown constitutive parameters. A subsequent difficulty is to build systems of equations which are well conditioned. Moreover, in these previous studies focused on nearly incompressible solids, the identification of compressibility parameters was avoided by selecting virtual fields which would filter out the effects of the hydrostatic pressure in the principle of virtual power. This was generally required as the volume changes could not be measured with enough accuracy by optical techniques, causing too much uncertainty on the local values of the hydrostatic pressure anyhow. Generating virtual fields that can filter out the hydrostatic pressure was never easy with analytic functions. The use of a finite-element implementation of the VFM simplified this task [50] but the novel approach presented in the current paper further generalizes this for any situation. There is a price to pay though as the approach proposed in the current paper is iterative, requiring to choose a first guess of the unknown material parameters and repeating the identification a number of time. Fortunately, we found that the convergence of this iterative approach was always quadratic.

It is important to emphasize that any kinematically admissible virtual displacement with a non-zero volume change is always an option to identify material properties of compressible materials with the VFM. However, only very specific virtual fields can ensure that the final system of equations is well conditioned and it can be cumbersome to determine these virtual fields by trial and errors. For complex geometries and when there are several parameters to identify, as in the ONH problem shown in this paper, it is convenient to use our novel VFM approach, which automatically generates the virtual fields as solutions of Eq. 42. Although these specific virtual fields satisfy the equilibrium conditions written in Eq. 42, virtual fields in general do not have to satisfy such equations.

To test the feasibility of the novel VFM approach, we first utilized the simplest hyperelastic model: the Neo-Hookean model. This model remains very popular for many biological tissues as for instance ocular tissues [23]. The novel VFM approach, which is iterative, showed a fast and quadratic convergence for neo-Hookean identification. Additionally, we employed the proposed approach for more sophisticated model, such as the Mooney-Rivlin model and the Veronda-Westmann model. The estimation results demonstrated that the proposed method is able to identify the hyperelastic parameters of such models, and even that constitutive equations involving an exponential could still be identified with high accuracy even using a single loading stage.

We then applied the proposed VFM method to the clinically-relevant challenge of extracting material properties of ONH tissues from full-field OCT data. Modeling the LC in living subjects is difficult, as this tissue is strongly inhomogeneous and exhibits very large variations in strain. In addition, OCT imaging does not allow the user to definitively probe the tissue itself and often cannot fully resolve tissue boundaries, complicating inverse finite element methods which rely upon strictly defined geometry to delineate tissues and material models. Results obtained on simulated data are very promising, showing that our VFM approach can remove these difficulties by providing material properties in different regions of the ONH. Future use on experimental measurements will allow to examine regional stiffness variations, which otherwise may have been neglected.

Despite these significant improvements of the VFM, the content of the current paper remains mostly theoretical, introducing the concepts, verifying them for simple cases and validating the approach on a first case study. The main limitation of the current approach is that it was developed in Matlab. It needs to be fully incorporated into a finite-element package to optimize computational costs and constitute a systematic alternative to FEMU techniques [10, 11] for all identification problems in hyperelasticity based on full-field displacement measurements.

Another limitation of the VFM is that it requires full-field data to achieve parameter identification. Although it performs very well when these full-field data are available, other methods based on the FEMU approach [10] do not have this limitation. For many years, full-field data were mostly available in two-dimensional bodies (membranes, shells) on which displacement fields could be measured using the DIC technique [22, 27, 31]. However, bulk measurements of displacements fields using digital volume correlation and imaging techniques such as Optical Coherence Tomography or Magnetic Resonance Imaging are becoming commonplace in soft tissues [23, 24, 25, 32, 33, 34, 35], offering more and more applications for the VFM in the biomedical field.

## 5 Conclusions

In this paper we presented a novel framework of the virtual fields method based on finite-element implementation and automatic generation of virtual fields. The proposed approach makes a great improvement in the theoretical aspects of the VFM as the choice of virtual fields is usually critical in the VFM. Future work will be focused on applying this finite-element based VFM into more complex cases with potential mechanobiological and clinical applications.

## 6 Acknowledgements

The authors acknowledge the support from the Fundamental Research Funds for the National Key Research and Development Plan (2016YFB0201601), National Natural Science Foundation (11732004, 11821202,12002075), Program for Changjiang Scholars, Innovative Research Team in University (PCSIRT), 111 Project (B14013), the Fundamental Research Funds for the Central Universities (Grant No. DUT19RC(3)017) in China and the European Research Council for grant ERC-2014-CoG BIOLOCHANICS.

